# HIPPOCAMPAL TAU-RELATED SIGNAL INDEXES HYPEREXCITABILITY AND NREM SLEEP–MEMORY DYSFUNCTION IN TEMPORAL LOBE EPILEPSY

**DOI:** 10.64898/2026.09.07.748279

**Authors:** Thaera Arafat, Katharina Schiller, Jack Lam, Nicolas Von Ellenrieder, Zhengchen Cai, Jordan DeKraker, Fatemeh Fadaie, Viviane Sziklas, Joelle Crane, Raúl R. Cruces, Robert Hopewell, Chris Hung-Hsin Hsiao, Gassan Massarweh, Jean-Paul Soucy, Kjell Erlandsson, Seyyed Ali Hosseini, Tevy Chan, Eliane Kobayashi, Pedro Rosa-Neto, Andrea Bernasconi, Neda Bernasconi, Matthias Koepp, Jean Gotman, Boris C. Bernhardt

## Abstract

**Background:** Sleep and memory disturbances are common in temporal lobe epilepsy (TLE), yet their relationship with tau-targeted neuroimaging measures remains unclear. We investigated whether *in-viv*o temporal ^18^F-MK6240 tracer retention relates to seizure burden, non–rapid eye movement (NREM) sleep microstructure, and memory performance in TLE.

**Methods:** This cross-sectional study was conducted from 2019 to 2024 at the Montreal Neurological Institute-Hospital. Eligible patients had unilateral TLE diagnosed according to International League Against Epilepsy criteria and underwent overnight electroencephalography (EEG) recordings, research-protocol 3T magnetic resonance imaging, and ^18^F-MK6240 positron emission tomography. Regional ^18^F-MK6240 standardized uptake value ratios (SUVRs), using the cerebellum as the reference region, were quantified in the ipsilateral hippocampus (defined a *priori* as the primary region of interest), as well as in peri-hippocampal and lateral temporal regions. Electroclinical and neuropsychological measures, including seizure burden (high vs low), N2 interictal epileptiform discharge frequency (frequent vs non-frequent), and memory performance (impaired vs intact), were classified according to predefined clinical criteria. N2 spindle rate and N3 slow-wave rate were quantified from overnight EEG recordings.

**Findings:** Of 70 eligible patients, 25 were included (eight [32%] females; mean age 35·2 years [SD 12·1]). Higher ipsilateral hippocampal ^18^F-MK6240 uptake was observed in participants with greater focal seizure burden (median [IQR] 0·70 [0·67–0·78] vs 0·64 [0·58–0·65], rank-biserial correlation=−0·58, 95% CI −0·82 to −0·19; p=0·02), was associated with lower N2 spindle rate (r=−0·42, 95% CI −0·70 to −0·03; p=0·04) and higher N3 slow-wave rate (r=0·47; 95% CI 0·09 to 0·73; p=0·02), and was also higher in participants with impaired than intact memory performance (n=21; mean [SD] 0·80 [0·12] vs 0·68 [0·10], Cohen’s d=1·15, 95% CI 0·07 to 2·21; p=0·04). Associations outside the hippocampus were generally less consistent and less precise, with the largest non-hippocampal estimate observed in peri-hippocampal regions.

**Interpretation:** Higher hippocampal ^18^F-MK6240 uptake was associated with greater focal seizure burden, altered NREM sleep microstructure, and impaired memory performance in TLE. These findings suggest that hippocampal ^18^F-MK6240 uptake may represent a candidate marker of circuit vulnerability associated with NREM sleep-memory dysfunction in TLE, warranting validation in larger longitudinal studies.

**Funding:** Canadian Institutes of Health Research, Fonds de la Recherche du Québec – Santé, Wellcome Trust, Natural Sciences and Engineering Research Council of Canada, Brain Canada, SickKids Foundation.

## Introduction

Sleep and memory disturbances are among the most frequent and disabling symptoms across neurological disorders characterized by neuronal hyperexcitability, including epilepsy (1, 2) and neurodegenerative disorders (3).

In neurodegenerative disorders, cortical accumulation of tau can be measured using positron emission tomography (PET) (4) with second-generation tau tracers such as ^18^F-MK6240, which demonstrate high specificity for neurofibrillary tangles and high sensitivity for medial temporal and neocortical tau deposition across the Alzheimer disease (AD) spectrum (5, 6). Studies have also demonstrated that higher tau biomarker levels are associated with alterations in sleep microstructure, including reduced non–rapid eye movement (NREM) sleep spindle density and slow-wave activity (3, 7). Although these relationships are increasingly recognised in AD, cognitive dysfunction in epilepsy emerges within a more complex clinical context shaped by the underlying epileptogenic pathology, neuropsychiatric and sleep comorbidities, and cumulative treatment effects. Converging evidence indicates that occurrence of epileptiform activity, both seizures and interictal epileptic discharges (IEDs), can disrupt sleep by inducing arousals, fragmenting sleep continuity, and altering microarchitecture (2). This disruption is particularly relevant during NREM sleep, a critical period for memory consolidation (8). Thus, epileptiform activity during NREM may compromise the physiological mechanisms required for memory stabilization.

Consistent with this framework, histological analyses of *ex-vivo* surgical specimens and *post-mortem* brain tissue provide growing evidence of hyperphosphorylated tau accumulation even in middle-aged adults with epilepsy (9). These alterations have been reported to be associated with neuronal damage, disruption of neural network function, and cognitive impairment, suggesting a potential mechanistic link between epilepsy and neurodegeneration (10). Complementing these findings, preliminary *in-vivo* PET studies using the ^18^F-MK6240 tracer have suggested that higher 18F-MK6240 signal in temporal lobe epilepsy (TLE) is associated with poorer executive and episodic memory performance (11). Recent *in-vivo* evidence using the ^18^F-Flortaucipir (AV-1451) tracer suggests that higher tracer signal is associated with clinical and electroencephalographic markers of epilepsy severity (12). Together, these observations provide preliminary evidence linking tau-targeted PET signal separately with epileptic activity and cognitive dysfunction in focal epilepsy; however, its relationship with network hyperexcitability, NREM sleep microstructure, and memory has not been examined within the same cohort.

Building on these findings, we hypothesised that greater ^18^F-MK6240 signal within the epileptogenic temporal lobe, particularly the hippocampus, would be associated with higher seizure burden and disrupted NREM sleep microstructure, which is critical for memory consolidation. To examine the interplay among these processes, we combined *in-vivo* ^18^F-MK6240 PET with electroclinical and behavioural measures in patients with TLE.

## Methods

### Study design

This cross-sectional observational study investigated associations between regional ^18^F-MK6240 uptake, electroclinical markers of hyperexcitability, and NREM sleep–memory measures in TLE. This study is reported in accordance with the Strengthening the Reporting of Observational Studies in Epidemiology (STROBE) guidelines.

### Participants

From 2019 to 2024, 70 adult patients with unilateral TLE were admitted to the Epilepsy Monitoring Unit (EMU), Montreal Neurological Institute-Hospital for comprehensive clinical evaluation. Diagnosis and classification were established according to International League Against Epilepsy criteria using long-term video-electroencephalography (EEG) monitoring, routine neuroimaging (1.5T or 3T, depending on clinical availability) and, when available, neuropsychological assessments. All underwent research-protocol 3T structural magnetic resonance imaging (MRI), which was used for all quantitative analyses, including hippocampal segmentation, region-of-interest definition, and PET co-registration.

Patients meeting inclusion criteria were enrolled in the study and underwent ^18^F-MK6240 PET imaging. Exclusion criteria included prior epilepsy surgery, chronic use of sleep medications, and a clinical diagnosis of a neurodegenerative disease.

All study participants provided written informed consent in accordance with the institutional Research Ethics Board (Montreal Neurological Institute and Hospital; #2018-4148).

### 3T MRI acquisition and processing

Structural MRI was acquired at the McConnell Brain Imaging Centre using a 3T Siemens Magnetom Prisma-Fit scanner with a 64-channel head coil. T1-weighted images were obtained using a 3D magnetization-prepared rapid gradient-echo sequence (MPRAGE; 0·8 mm isotropic voxels, TR 3200 ms, TE 3·14 ms, TI 900 ms, flip angle 9°). T1-weighted images were processed using micapipe v0·2·3 (13) (https://micapipe.readthedocs.io/) in combination with FreeSurfer v6·0 (14) (https://surfer.nmr.mgh.harvard.edu/) to perform cortical surface reconstruction and segmentation. All segmentations were manually inspected and corrected, and each T1-weighted MRI was deobliqued and reoriented to LPI orientation (left-right, posterior-anterior, inferior-superior).

Hippocampal structures were subsequently mapped to a standardized unfolded hippocampal surface using HippUnfold v1·3·0 (15). Hippocampal volumes were derived from 3T T1-weighted MRI using a validated automated segmentation pipeline. Volumes were corrected for intracranial volume and converted to age-adjusted z-scores using an age-matched healthy control cohort acquired with the same imaging protocol (15-17). Hippocampal atrophy was defined a priori as MRI-derived hippocampal volume or asymmetry indices exceeding 1·5 standard deviations relative to the corresponding age-matched healthy control distribution (16, 18). Quantitative classifications were cross-checked against the corresponding clinical neuroradiology reports (**see Supplementary Methods for details**).

### 18F-MK6240 PET acquisition and processing

Tau-PET imaging with ^18^F-MK6240, a highly selective radiotracer for neurofibrillary tangles (19), was performed on a Siemens HRRT scanner. The radiotracer was synthesized at the PET Unit of the Montreal Neurological Institute following published protocols (20). Images were acquired 90– 110 minutes after intravenous injection, with a mean injected dose of 224 MBq (SD = 19 MBq).

PET data were reconstructed and subsequently processed using a standardized in-house pipeline previously described in published work from our group (11, 21). Reconstruction used an ordered-subset expectation maximization algorithm. A short transmission scan (11 min) with a rotating 137Cs source was used for attenuation correction during reconstruction. Reconstruction also incorporated correction for dead time, decay, random coincidences, and scattered coincidences. Four dynamic^18^F-MK6240 frames were co-registered and averaged to generate a static PET image.

Subsequent image processing included registration of averaged ^18^F-MK6240 PET volume to the participant’s T1-weighted MRI using an affine registration based on cross-correlation and histogram matching (11), followed by projection onto each participant’s cortical midthickness surface. No post-reconstruction partial-volume correction. As a sensitivity analysis, the principal hippocampal analyses were repeated using iterative Yang partial-volume correction; details are provided in the **Supplementary Methods**. All participants were processed using the same reconstruction and analysis pipeline. Surface-based ^18^F-MK6240 maps were subsequently smoothed using a 10-mm FWHM surface-based kernel and resampled to the fsLR-32k standard surface. Standardized uptake value ratio (SUVR) maps were subsequently calculated using cerebellar grey matter as the reference region and served as the primary quantitative PET outcome.

Region of interest (ROI) analyses were conducted using FreeSurfer v6·0, with mean SUVR values extracted from anatomically defined volumetric ROIs in the epileptogenic temporal regions (**Figure 1A**). The primary ROI was the ipsilateral hippocampus, selected based on our *a priori* hypothesis that hippocampal ^18^F-MK6240 signal would be associated with sleep and memory impairment in TLE. To contextualise hippocampal uptake alterations in a broader anatomical context within the temporal lobe, we additionally quantified ^18^F-MK6240 SUVR across peri-hippocampal regions (amygdala, parahippocampal gyrus, entorhinal cortex, and temporal pole), and lateral temporal regions (superior, middle, and inferior temporal gyri and fusiform gyrus).

**Figure 1.**
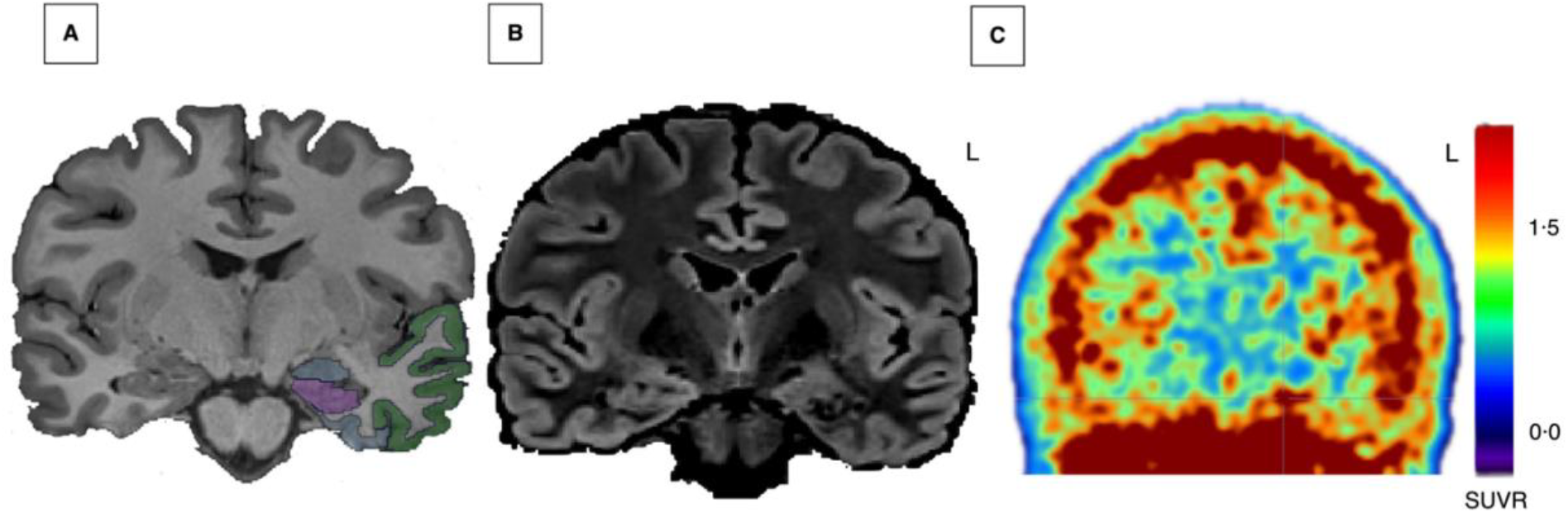
^18^F-MK6240 PET and region of interest definition (ROI). (A) T1-weighted 3T MRI illustrating ROIs in the ipsilateral temporal lobe: hippocampus (*magenta*); peri-hippocampal regions (amygdala, parahippocampal gyrus, entorhinal cortex, temporal pole; *blue*), and lateral temporal regions (superior, middle, inferior temporal gyri, fusiform gyrus; *green*). (B) Structural research 3T MRI (FLAIR sequence) from a 28-year-old man with late-onset left temporal lobe epilepsy, demonstrating a cavernoma within the left hippocampus; this image is shown for clinical illustration and was not used for PET co-registration. (C) ^18^F-MK6240 PET image co-registered to the corresponding T1-weighted 3T MRI from the same participant, showing bilateral neocortical ^18^F-MK6240 SUVR uptake with left-sided predominance, particularly within temporal lobe. Regional SUVR values were higher on the left than the right for the peri-hippocampal region (1·00 vs 0·86), lateral temporal region (1·21 vs 1·17), and hippocampus (0·92 vs 0·65). PET images correspond to the standard processing pipeline used for all quantitative analyses and are displayed using a continuous SUVR colour scale (0·0–1·5), without statistical thresholding or masking.

### Seizure burden scoring

Seizure burden was assessed based on seizure type and frequency obtained from retrospective clinical history documented during EMU admission. Patients were classified a priori into high and low seizure burden groups using clinically defined thresholds based on seizure frequency and type, which have previously been linked to cognitive outcomes in TLE (22-24). High seizure burden was defined as a focal to bilateral tonic–clonic seizures (FBTCs) ≥3 per year or focal seizures occurring weekly or more frequently, whereas low seizure burden was defined as fewer than three FBTCs per year or focal seizures occurring less than weekly. The prespecified primary analyses used clinically defined seizure-burden categories to facilitate clinical interpretation, whereas continuous focal seizure-frequency analyses were performed as complementary sensitivity analyses to assess the robustness of the findings to the method of exposure specification (see **Supplementary Methods** for details).

### Overnight scalp EEG recordings

Scalp EEGs were recorded during the second night of admission to the EMU, from 20:00 to 08:00, while patients remained on the same dosage of antiseizure medications and had adapted to the new environment. Overnight EEG recordings were acquired using the Nihon Koden system (Tokyo, Japan) with 19 glued electrodes placed according to the international 10–20 EEG system, supplemented by two orbitofrontal (SO1/SO2), two zygomatic (Zy1/Zy2), and six inferior temporal chain electrodes (F9/F10, T9/T10, P9/P10); signals were sampled at 1000 Hz and referenced to Cz.

### Sleep scoring and automatic detection of NREM sleep spindles and slow-wave events

All EEG recordings were manually scored by a sleep expert (K.S.) in the Stellate Reviewer software (Montreal, Canada), in accordance with the American Academy of Sleep Medicine criteria, in 30-second epochs and using a bipolar montage (25), and were performed blinded to imaging and clinical outcome variables. For automated event detection, all N2 and N3 sleep epochs were selected. EEG channels exhibiting continuous artifacts were excluded from further analysis. Sleep spindles and slow waves were automatically detected using a validated algorithm (26) applied to the average reference montage. Spindles were defined by a frequency range of 10–16 Hz and a duration of 0·5–3 s, while slow waves were identified within a frequency range of 0·5– 4 Hz. High-amplitude N3 slow waves detected in our analysis may reflect transient increases in network excitability, as slow-wave dynamics are known to modulate cortical states and facilitate epileptic activity (27). Global spindle and slow-wave rates were calculated as the mean number of events per minute, averaged across all channels for each patient.

### IEDs marking and frequency index

IEDs, including isolated spikes (low, medium, or high amplitude), spike–slow wave, and bursts of epileptiform activity lasting less than 10 sec, were manually marked by a board-certified epileptologist (T.A.) using Stellate Reviewer software (Montreal, Canada), with marking performed blinded to imaging and clinical data. Analyses were performed on the first 30 min of consecutive 30 sec epochs of stage N2 sleep using bipolar and average montages. Isolated interictal discharges were annotated at their peak across all channels, whereas bursts were annotated by marking onset and offset. All interictal epileptiform abnormalities were analysed and expressed as events per minute of N2 sleep. IED rates were classified as frequent (≥1 event/min) and non-frequent (occasional/rare, <1 event/min) groups, consistent with standardized EEG frequency descriptors (28).

### Memory assessment

Verbal and nonverbal learning and memory scores were obtained from standardized neuropsychological assessments conducted by certified neuropsychologists (V.S. and J.C.). Data were available for 21 of 25 patients and were included in analyses. The median interval between the memory testing and tau-PET imaging was 5·1 months (IQR: 1·0–20·6 months). Verbal memory was assessed by either the Rey Auditory Verbal Learning Test (RAVLT) (29), and/or the Nicole Abstract Verbal Learning Test (NAVLT) (30). Both tests comprise learning 15 words over 5 trials (T1–T5), followed by exposure to and immediate recall of a second list of words (interference list), post-interference recall of the first list (T6), and a final delayed recall of the first list after 30 minutes (T7). The RAVLT and NAVLT are administered in a similar way and considered analogous tests, with the differences being the concreteness/abstractness of the words and that the RAVLT is given orally, while NAVLT is presented visually, and responses are written. Nonverbal learning was assessed using the Aggie Figures Learning Test (AFLT) (31). The test consists of five learning trials of abstract figures (T1–T5), followed by an interference trial (second set of figures), post-interference (T6) and 30 min delayed (T7) recalls of the first set of figures.

To derive an overall index of memory performance, we computed a composite by averaging age- and sex-adjusted z-scores from verbal and nonverbal total learning measures, based on published normative data for the RAVLT (32), and AFLT (33). The composite z-score was computed as the mean of the verbal and nonverbal total learning z-scores for each patient. Participants were classified as impaired if their memory composite z-score was below −1·3, corresponding to approximately the 10th percentile and consistent with established neuropsychological definitions of clinically meaningful cognitive impairment (34-36), and as intact otherwise. Primary analyses used this clinically defined classification to identify participants with clinically meaningful memory impairment, whereas continuous analyses of the memory score were performed as sensitivity analyses.

### Statistics

Primary analyses were hypothesis-driven and prespecified. Sensitivity analyses were performed to assess the robustness of the primary findings.

Primary statistical analyses focused on the ipsilateral hippocampus, with peri-hippocampal and lateral temporal regions examined as prespecified regional comparisons using extracted ^18^F-MK6240 SUVR values. Continuous variables were summarised as mean (SD) or median (IQR), as appropriate.

Associations between regional ^18^F-MK6240 SUVR and continuous measures of sleep microstructure (N2 spindle and N3 slow-wave rates) were assessed using Pearson correlation analyses. Group differences in regional ^18^F-MK6240 SUVR were evaluated for prespecified clinically defined variables, including seizure burden (high versus low), IED rate (frequent versus non-frequent), and memory performance (impaired versus intact). Distributional assumptions were assessed using the Shapiro–Wilk test and Q–Q plots, and scatterplots were examined for linearity for correlation analyses. Between-group comparisons were performed using independent-samples t-tests or Mann–Whitney U tests, as appropriate. For analyses additionally examined using linear regression, linearity, residual normality, and homoscedasticity were assessed using the relevant diagnostic plots, and leverage and Cook’s distance were examined to identify potentially influential observations. All observations were retained in the primary analyses because no prespecified clinical or methodological criterion for exclusion was met. Sensitivity analyses excluding observations with relatively high leverage were performed where relevant.

Sensitivity analyses included (i) adjustment for hippocampal volume in models examining the association between ^18^F-MK6240 uptake and memory performance; (ii) a multivariable logistic regression analysis of memory impairment (impaired vs intact), including seizure focus laterality as a clinically relevant covariate, given its established association with memory performance in TLE (37); (iii) assessment of anti-seizure medication (ASM) burden in relation to memory performance and sleep microstructure; (iv) analyses treating focal seizure frequency and memory performance as continuous measures; and (v) analysis using standardised regional ^18^F-MK6240 z-scores derived from an age-matched healthy control cohort; and (vi) repetition of the principal hippocampal analyses using iterative Yang partial-volume-corrected ^18^F-MK6240 SUVR values. Sensitivity analyses were performed to assess the robustness of the primary findings and were interpreted according to the magnitude, direction, and precision of the estimated effects, rather than P values alone **(see Supplementary Methods)**.

Biological sex was included as a covariate in prespecified sensitivity analyses where appropriate. Sex-stratified analyses were not performed because of limited statistical power (**Table 1**).

**Table 1.** Demographic and Clinical Characteristics of Participants. **Abbreviations:** ASM=anti-seizure medication; BRV=brivaracetam; CBZ=carbamazepine; CLB=clobazam; CZP=clonazepam; ESL=eslicarbazepine; FBTC=focal to bilateral tonic-clonic seizure; IED=interictal epileptiform discharge; IQR=interquartile range; LCM=lacosamide; LEV=levetiracetam; LTG=lamotrigine; MTS=mesial temporal sclerosis; NREM=non-rapid eye movement; N2/N3=NREM sleep stages N2 and N3; PER=perampanel; PHT=phenytoin; SD=standard deviation; TLE=temporal lobe epilepsy; TPM=topiramate; VPA=valproate. **Note:** Data are presented as mean (SD; range) or median (IQR; range) unless otherwise indicated. Patients may contribute to more than one ASM class due to polytherapy.

| Characteristic | Values |
| --- | --- |
| **Demographics** |  |
| Age, years, mean (SD; range) | 35.2 (12.1; 23–65) |
| Sex, n (%) | Female 8 (32%); Male 17 (68%) |
| Race/ethnicity, n (%) | White 13 (52%); Black/African descent 5 (20%); Hispanic/Latino 3 (12%); Middle Eastern 1 (4%); Other/uncertain 3 (12%) |
| Handedness, n (%) | Left-handed 4 (16%); Right-handed 21 (84%) |
| Education, years, mean (SD; range) | 13.5 (2.1; 9–18) |
| **Epilepsy characteristics** |  |
| Age at epilepsy onset, mean (SD; range) | 21.8 (14.8; 1–60) |
| Disease duration, years, mean (SD; range) | 12.8 (8.3; 2–30) |
| Epilepsy type, n (%) | TLE 21 (84%); TLE+ 4 (16%) (3 Fronto-temporal; 1 Temporo-insular) |
| Focus lateralization | Left 13 (52%); Right 12 (48%) |
| Focal seizure frequency, median (IQR; range), per week | 1 (0.25–2.25; 0–14) |
| **FBTC seizure frequency, median (IQR; range), per year** | 1 (0–3; 0–26) |
| **MRI lesion category (n, %)** | Non-lesional 16 (64%); MTS 5 (20%); Other lesions 4 (16%) |
| **EEG metrics** |  |
| **IED rate during N2 sleep, median (IQR; range), per min** | 0.05 (0.01–0.11; 0–0.82) |
| **N2 spindle rate, median (IQR; range), per min** | 3.87 (2.97–4.68; 2.21–6.54) |
| **N3 slow-wave rate, median (IQR; range), per min** | 2.95 (0.50–5.90; 0–12.3) |
| **Cognitive measures** |  |
| **Memory performance (z-score), median (IQR; range)** | –0.24 (–0.68–0.17; –1.93–0.98) |
| **ASMs** |  |
| **Number of ASMs, median (IQR)** | 2 (2–3) |
| **ASM class, n (%)** | Sodium channel blockers (CBZ, LTG, LCM, ESL, PHT, TPM) 20 (80%); SV2A modulators (LEV, BRV) 10 (40%); GABAergic agents (CLB, CZP, VPA) 9 (36%); AMPA antagonist (PER) 3 (12%) |

Effect estimates are reported according to analysis type, including Pearson correlation coefficients (r), Cohen’s d for parametric group comparisons, rank-biserial correlations for Mann–Whitney U comparisons, and regression coefficients where applicable, together with corresponding 95% confidence intervals and two-sided P values. Because the primary analyses were prespecified and hypothesis-driven, no adjustment for multiple comparisons was applied. All statistical analyses were performed using JASP (v0·95·4) and R (v4·4·1).

## Results

Of 70 eligible patients, five (7%) were excluded because of previous epilepsy surgery and 40 (57%) declined ^18^F-MK6240 PET imaging, leaving a final cohort comprised 25 (36%) patients (eight [32%] females; mean [SD] age 35·2 [12·1] years; mean [SD] epilepsy duration 12·8 [8·3] years). The study recruitment and inclusion process are summarised in **Supplementary Figure 1**. Among the 25 participants, 21 patients (84%) had classical TLE, and four (16%) had TLE+ (fronto-temporal epilepsy, n=3; temporo-insular epilepsy, n=1). Seven of 25 participants (28%) met the predefined criterion for hippocampal atrophy based on MRI-derived hippocampal volume or asymmetry indices (**see Supplementary Methods**).

Of the 25 participants, five (20%) underwent temporal lobe surgery following study participation, including selective amygdalohippocampectomy (n=2), corticoamygdalohippocampectomy (n=2), and focal lesionectomy (n=1). At 1-year follow-up, four patients achieved modified Engel (38) Class IA seizure freedom and Class IVB in one.

Histopathological diagnosis according to International League Against Epilepsy criteria (39) was extracted from clinical pathology reports. Three cases (60%) showed double or dual pathology (two with focal cortical dysplasia (FCD) type IIA and hippocampal gliosis, and one with FCD IIA and hippocampal sclerosis type 1), whereas two cases (40%) demonstrated a single pathology (collagenized hippocampal cavernoma in one and mild cortical dysplasia in another).

To characterize tau pathology in available surgical tissue, immunohistochemistry for phosphorylated tau (pTau) (AT8; recognizing pTau at Ser202/Thr205; 1:1200; Thermo Fisher Scientific, MN1020) was performed on 12 tissue blocks from five participants using established protocols. Tau immunoreactivity was qualitatively classified as absent (no detectable tau), subtle (weak subpial staining or rare positive neurons), or strong (robust intraneuronal positivity or tufted-like staining). Consistent with prior studies (40, 41), tau accumulation was most pronounced in a hippocampal block surrounding a collagenized cavernoma, showing intracytoplasmic neuronal positivity with tufted-like staining in adjacent regions. The participant with an MRI-defined left hippocampal cavernoma (**Figure 1B**) also demonstrated high hippocampal ^18^F-MK6240 signal compared with the remainder of the cohort (**Figure 1C)**.

Blocks from hippocampal gliosis ranged from absent to subtle, whereas FCD type IIA blocks ranged from absent to subtle (mild subpial positivity observed in one case). Mild cortical dysplasia blocks were negative. Participant demographic and clinical characteristics are summarised in **Table1**.

### Ipsilateral hippocampal^18^F-MK6240 uptake: associations with electroclinical and behavioural measures

Higher hippocampal ^18^F-MK6240 uptake was observed in participants with higher focal seizure burden, altered NREM sleep microstructure, and impaired memory performance.

Patients with weekly or more frequent focal seizures (n=16) showed higher tracer uptake than those with less frequent focal seizures (n=9) (median [IQR] 0·70 [0·67–0·78] vs 0·64 [0·58–0·65]; rank-biserial correlation=−0·58, 95% CI −0·82 to −0·19; p=0·02; **Figure 2A**), whereas the estimated difference between patients with ≥3 FBTCs/year (n=8) and those with fewer FBTCs (n=17) was smaller and imprecise (mean [SD] 0·73 [0·14] vs 0·69 [0·09]; Cohen’s d=0·38, 95% CI −0·47 to 1·23; p=0·38; **Figure 2B**). Similarly, the estimated difference between patients with frequent IEDs (≥1 event/min; n=15) and those with non-frequent IEDs (n=10) was moderate but imprecise (mean [SD] 0·73 [0·11] vs 0·66 [0·11]; Cohen’s d=0·58, 95% CI −0·25 to 1·39; p=0·17; **Figure 2C**).

**Figure 2.**
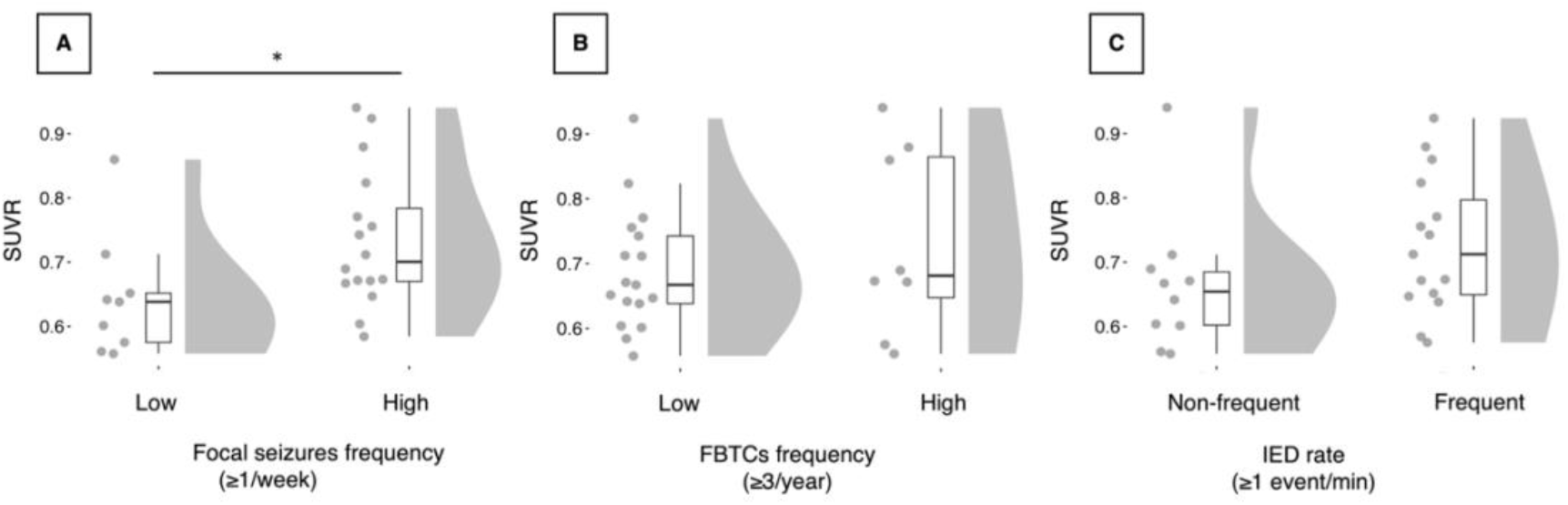
Ipsilateral hippocampal ^18^F-MK6240 uptake and seizure burden measures. (A) Higher ^18^F-MK6240 uptake was observed in the high focal seizure burden group (≥1/week; n=16) compared with the low-burden group (n=9) (median [IQR] 0·70 [0·67–0·78] vs 0·64 [0·58–0·65]; rank-biserial correlation=−0·58, 95% CI −0·82 to −0·19; p=0·02). (B) No clear evidence of a difference in tracer uptake were observed between focal to bilateral tonic–clonic seizure burden (FBTCs; ≥3/year; n=8 vs n=17) (mean [SD] 0·73 [0·14] vs 0·69 [0·09]; Cohen’s d=0·38, 95% CI −0·47 to 1·23; p=0·38). (C) No clear evidence of a difference in tracer uptake were observed between interictal epileptiform discharge (IEDs; ≥1 event/min; n=15 vs n=10) (mean [SD] 0·73 [0·11] vs 0·66 [0·11]; Cohen’s d=0·58, 95% CI −0·25 to 1·39; p=0·17).

Higher hippocampal ^18^F-MK6240 uptake was inversely associated with N2 spindle rate (n=25; r=−0·42, 95% CI −0·70 to −0·03; p=0·04; **Figure 3A**) and positively associated with N3 slow-wave rate (n=25; r=0·47, 95% CI 0·09 to 0·73; p=0·02; **Figure 3B**). In a sensitivity analysis excluding the two observations with relatively high leverage, the association between hippocampal ^18^F-MK6240 uptake and N3 slow-wave rate remained positive (n=23; r=0·38, 95% CI −0·04 to 0·68; p=0·07; **Supplementary Table 7; Supplementary Figures 5 and 6**). Patients with impaired memory performance (n=5) showed higher tracer uptake than those with intact performance (n=16) (mean [SD] 0·80 [0·12] vs 0·68 [0·10]; Cohen’s d=1·15, 95% CI 0·07 to 2·21; p=0·04; **Figure 3C**).

**Figure 3.**
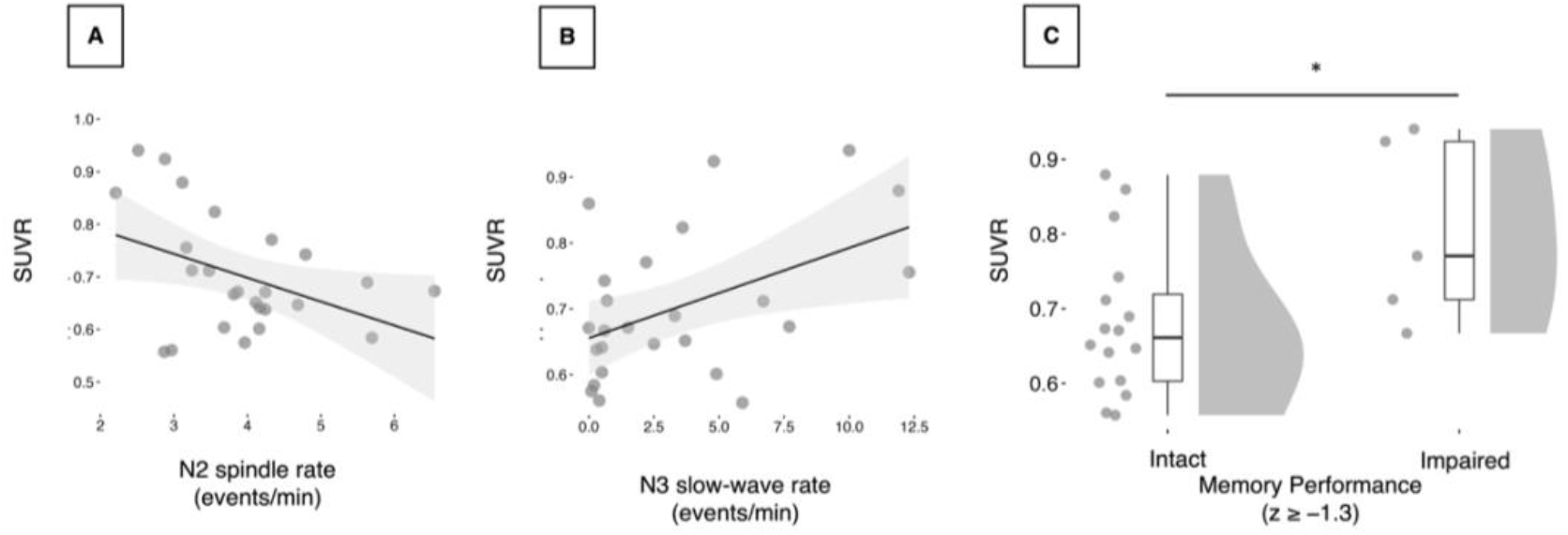
Ipsilateral hippocampal ^18^F-MK6240 uptake, NREM sleep microstructure and memory performance. (A) Higher ^18^F-MK6240 uptake was associated with reduced N2 spindle rate (n=25; r=−0·42, 95% CI −0·70 to −0·03; p=0·04). (B) Higher ^18^F-MK6240 uptake was associated with elevated N3 slow-wave rate (n=25; r=0·47, 95% CI 0·09 to 0·73; p=0·02). (C) Patients with impaired memory performance (n=5) showed higher ^18^F-MK6240 uptake than those with intact performance (n=16) (mean [SD] 0·80 [0·12] vs 0·68 [0·10]; Cohen’s d=1·15, 95% CI 0·07 to 2·21; p=0·04).

To further assess the robustness of these findings, we undertook several prespecified sensitivity analyses.

The association between hippocampal ^18^F-MK6240 uptake and memory impairment was directionally preserved after adjustment for hippocampal volume and after accounting for seizure focus laterality, although estimates were less precise after adjustment for hippocampal volume **(Supplementary Table 1**).

ASM burden did not differ between participants with impaired and intact memory performance (n=21; Cohen’s d=−0·33, 95% CI −1·33 to 0·69; p=0·53). ASM burden was positively associated with N2 spindle rate (n=25; r=0·65, 95% CI 0·34 to 0·83; p<0·001), but not with N3 slow-wave rate. Adjustment for ASM burden yielded an association in the same direction, although the estimate was less precise (n=25; B=−0·05, 95% CI −0·11 to 0·01; p=0·08; **Supplementary Table 2**).

When seizure burden and memory performance were analysed as continuous variables rather than predefined groups, the estimated associations with hippocampal ^18^F-MK6240 SUVR were directionally consistent with the primary analyses but were small and imprecise (focal seizure burden: n=25; r=0·22, 95% CI −0·19 to 0·57; p=0·28; memory performance: n=21; r=−0·21, 95% CI −0·59 to 0·24; p=0·35; **Supplementary Table 3**).

The direction and overall pattern of the principal hippocampal associations also remained consistent after iterative Yang partial-volume correction (**Supplementary Table 6**).

### Beyond the ipsilateral hippocampus: weaker ^18^F-MK6240 uptake associations

To contextualize hippocampal findings within the broader temporal lobe, ^18^F-MK6240 uptake was quantified across peri-hippocampal and lateral temporal regions in the epileptogenic hemisphere (see Methods for ROI definitions).

Overall, associations between extra-hippocampal ^18^F-MK6240 uptake and electroclinical or behavioural measures were generally weaker and less consistent than those observed for the hippocampus, suggesting a relative hippocampal predominance of the observed pattern.

Peri-hippocampal ^18^F-MK6240 uptake was higher in patients with higher focal seizure burden (n=16) compared with the low-burden group (n=9) (mean [SD] 0·83 [0·09] vs 0·76 [0·07]; Cohen’s d= 0·79, 95% CI, −0·07 to 1·63; p=0·07). Estimated associations with FBTC seizure burden, IED frequency, NREM sleep microstructure, and memory performance were small and imprecise. In lateral temporal regions, estimated associations with electroclinical and memory measures were generally small and imprecise. Detailed results are shown in **Supplementary Figures 3 and 4**.

Sensitivity analyses using age-normalised ^18^F-MK6240 z-scores derived from an age-matched healthy control cohort showed similar overall patterns of association across the prespecified ROIs (**Supplementary Table 4**).

## Discussion

This cross-sectional multimodal study supports the role of the hippocampus as a hub linking hyperexcitability, NREM sleep disruption, and ^18^F-MK6240 uptake-related cognitive vulnerability in TLE. Leveraging TLE as a human model of network hyperexcitability, we integrated *in-vivo* tau PET with 3T MRI, alongside electroclinical measures of seizure burden, NREM sleep, and memory. Given the central role of the hippocampus in the pathophysiology of TLE, as well as its key involvement in hippocampal–neocortical communication during sleep and memory processing, the ipsilateral hippocampus was defined *a priori* as the primary ROI. To further evaluate the relative regional specificity of our findings to the hippocampus, we examined effects in peri-hippocampal regions (*i*.*e*., the amygdala, parahippocampal gyrus, entorhinal cortex, and temporal pole) as well as lateral temporal regions (namely the superior, middle, and inferior temporal gyri, and fusiform gyrus).

The association between ipsilateral hippocampal ^18^F-MK6240 uptake and greater focal seizure burden is consistent with previous literature linking recurrent focal seizures to cognitive decline, whereas interictal or subclinical epileptiform discharges may produce transient cognitive deficits and contribute to gradual decline over time (1). In contrast, the estimated association between IED burden and hippocampal ^18^F-MK6240 uptake was weaker and less precise in this cohort, suggesting that the observed association may not extend uniformly across different measures of epileptic activity. Cerebrospinal fluid biomarker studies have also demonstrated elevated pTau in patients with focal or convulsive seizures, reflecting activation of neurodegenerative pathways in epilepsy (42). Consistently, patients with dementia with epileptiform discharges experience accelerated cognitive decline (43). Beyond seizure burden, the association between ^18^F-MK6240 uptake and NREM sleep microstructure, particularly N2 sleep spindles and N3 slow waves, is relevant to the role of hippocampal–neocortical networks in memory consolidation (44). Importantly, overnight EEG recordings were acquired while patients remained on stable ASM regimens, limiting within-subject medication variability during sleep measurements. Although ASM burden was associated with N2 spindle rate in our cohort, adjustment for ASM burden did not materially alter the direction of the association between N2 spindle rate and hippocampal ^18^F-MK6240 uptake. Nevertheless, given the heterogeneous effects of individual ASMs on sleep physiology, residual medication-related effects cannot be excluded (45). In epilepsy, NREM IEDs may disrupt these rhythms, perturb hippocampal–frontal networks, and contribute to cognitive impairment, and high-amplitude N3 slow waves detected in our analysis may transiently increase network excitability and facilitate epileptic activity (27, 46). Converging evidence from aging and neurodegenerative disorders further demonstrates that sleep spindle and slow-wave alterations co-occur with tau pathology; here, reduced spindle density has been associated with elevated CSF total and phosphorylated tau (7). These observations point to a potential convergence of seizure burden, NREM sleep alteration, and tau tracer retention within the hippocampus, raising the possibility of a mechanistic link between network hyperexcitability, sleep-memory dysfunction and tau-related processes in epilepsy. In this context, hippocampal ^18^F-MK6240 uptake–memory associations may reflect broader hippocampal memory function rather than lateralized, domain-specific patterns, as analyses incorporated both verbal and non-verbal measures, and accounted for focus laterality. These findings align with emerging work on the bidirectional relationship between epilepsy and dementia, which links pTau to cognitive dysfunction (10, 47). Beyond epilepsy, tau-PET studies demonstrate that baseline hippocampal tau predicts longitudinal memory decline and progression to dementia in individuals with mild cognitive impairment (48), supporting the potential relevance of hippocampal tau-related PET signal as a marker of cognitive vulnerability.

Beyond the ipsilateral hippocampus, estimated associations between ^18^F-MK6240 uptake and electroclinical measures were generally smaller or less consistent, with the clearest non-hippocampal finding being an association between peri-hippocampal tracer retention and higher focal seizure burden; associations in lateral temporal regions were generally small and imprecise. Taken together, the observed pattern of associations in the hippocampus suggests a relative hippocampal predominance of the electroclinical associations. This pattern raises the possibility of a hippocampus-centred vulnerability profile in TLE linking ^18^F-MK6240 tracer retention with measures of network hyperexcitability and NREM sleep–memory dysfunction (49).

Our multimodal approach provides novel insights into the potential biological and circuit-level processes underlying hippocampal MK6240 uptake–electroclinical associations in TLE, a human model of network hyperexcitability, supporting hippocampal ^18^F-MK6240 signal as a potential biomarker of circuit-level dysfunction and NREM sleep–cognitive impairment. These observations were derived from a deeply phenotyped patient cohort that underwent comprehensive clinical phenotyping and multimodal research investigations, including ^18^F-MK6240 PET, 3T structural MRI, overnight scalp EEG recordings, and standardized neuropsychological assessments. Although this approach yielded a uniquely integrated dataset, the modest sample size may have limited the precision of effect estimates, and the detection of some regional associations. Nevertheless, the primary hippocampal associations showed generally moderate-to-large effect estimates, supporting the potential clinical relevance of hippocampal ^18^F-MK6240 signal as a marker of regional vulnerability and circuit-level dysfunction in TLE, while requiring confirmation in larger cohorts.

Although tracer quantification and signal fidelity can vary across mesiotemporal and lateral temporal regions (21), the observed pattern of hippocampal associations was clinically relevant and consistent with established models of TLE. No post-reconstruction partial-volume correction was applied in the primary analyses. Although residual partial-volume effects cannot be excluded, particularly within mesiotemporal structures, the direction and overall pattern of the principal hippocampal findings were preserved in a sensitivity analysis using iterative Yang partial-volume correction. Validation using higher-resolution imaging approaches and multimodal methods will be required to further clarify the biological basis of these patterns. Importantly, although ^18^F-MK6240 is widely used as a marker of tauopathy in Alzheimer’s disease, the molecular specificity of its retention in TLE requires further validation.

Several sensitivity analyses assessed the robustness of the principal associations to measurable structural and clinical factors. The direction of the hippocampal ^18^F-MK6240 uptake–memory association was preserved after adjustment for hippocampal volume and seizure focus laterality, and the association between N2 spindle rate and hippocampal ^18^F-MK6240 uptake was not materially altered after adjustment for ASM burden. Similar overall patterns were also observed using age-normalised ^18^F-MK6240 z-scores. In complementary analyses treating seizure frequency and memory performance as continuous measures, effect estimates were generally smaller or less precise, with wide confidence intervals. These findings illustrate that the clinically defined and continuous analyses address complementary questions and should be interpreted together. Overall, these observations highlight the sensitivity of estimates to the specification of clinically defined versus continuous measures and support replication in larger cohorts using prespecified continuous and clinically meaningful outcomes. Residual confounding cannot be excluded given the modest sample size and observational design.

The cohort primarily comprised young adults with TLE, providing a relatively homogeneous population for mechanistic interpretation. This enhances interpretability within this well-defined group but may limit generalizability to older adults, female-dominated cohorts, or other epilepsy syndromes. The cross-sectional design also limits assessment of the temporal relationships among these electroclinical measures. Together, these observations highlight the value of the current cohort for advancing mechanistic insight and underscore the importance of future multi-center and longitudinal studies to extend these findings to more diverse epilepsy populations, clarify causal relationships, and evaluate the potential predictive value of hippocampal ^18^F-MK6240 signal for cognitive outcomes.

## Contributors

T.A.: conceptualisation, study design, patient recruitment, data acquisition and curation, EEG marking, formal analysis, project administration, and writing – original draft. K.S.: sleep EEG scoring, study design input, and writing – review and editing. J.L.: PET data processing and quantification, data extraction and validation, and writing – review and editing. N.V.E.: software development and methodology for automated sleep microstructure analysis, and writing – review and editing. J.D.: hippocampal image processing, data extraction, normalisation, and writing– review and editing. Z.C.: formal analysis and validation. F.F.: structural imaging extraction, normalisation, and writing–review and editing. V.S. and J.C.: neuropsychological data acquisition and writing – review and editing. R.R.C.: PET data processing. R.H., C.H.-H.H., and G.M.: PET acquisition coordination, data acquisition, and writing–review and editing. J.P.S.: writing – review and editing. K.E.: quality assurance and correction of imaging code. T.C. and S.A.H.: methodology, data validation, and writing – review and editing. E.K.: conceptualisation, study design. P.R.-N.: resources, methodology, and writing – review and editing. A.B. and N.B.: resources and funding acquisition. M.K.: conceptualisation and writing – review and editing. J.G.: conceptualisation, supervision, study design, EEG analysis guidance, and writing – review and editing. B.C.B.: conceptualisation, supervision, study design oversight, resources, funding acquisition, and writing – review and editing. T.A. and B.C.B. had full access to all the data in the study and had final responsibility for the decision to submit for publication. All authors reviewed and approved the final manuscript.

## Data sharing

De-identified derived data will be made available on the Open Science Framework upon publication of this Article.

## Acknowledgments

We thank the patients and healthy volunteers who participated in this study. We acknowledge the McConnell Brain Imaging Centre (BIC) PET and MRI units at the Montreal Neurological Institute N(MNI) for technical support, and the clinical neurophysiology and epilepsy teams for facilitating access to EEG recordings and clinical data. We are grateful to Dr Raluca Pana, Dr François Dubeau, Dr Martin Veilleux, and Dr Andrea Bernasconi for their support of the clinical research environment.

T.A. and F.F. are supported by a Savoy Foundation Scholarship. J.L. is funded by the Fonds de la Recherche du Québec – Santé (FRQS) and the Ministère de la Santé et des Services sociaux du Québec (MSSS). P.R.-N. and R.R.C. are supported by FRQ-S, the MNI Jeanne Timmins Costello Fellowship, and the Healthy Brains, Healthy Lives Entrepreneur Postdoctoral Fellowship. M.K. is supported by the Wellcome Trust (221934/Z/20/Z). B.C.B. acknowledges support from the Canadian Institutes of Health Research (CIHR; FDN-154298, PJT-174995, PJT-206196, CIHR PJT-203761, CIHR PJT-191853), SickKids Foundation (NI17-039), Natural Sciences and Engineering Research Council (NSERC RGPIN-2025-05932), Azrieli Center for Autism Research of the Montreal Neurological Institute (ACAR), BrainCanada, FRQ-S, the Helmholtz International BigBrain Analytics and Learning Laboratory (Hiball), Healthy Brains and Healthy Lives (HBHL), Centre for Aging + Brain Health Innovation (CABHI), the Canada Research Chairs Program (CRC), and the Centre of Excellence in Epilepsy at the Neuro (CEEN).

