## Supplementary Material for "HIPPOCAMPAL TAU-RELATED SIGNAL INDEXES HYPEREXCITABILITY AND NREM SLEEP–MEMORY DYSFUNCTION IN TEMPORAL LOBE EPILEPSY"

#### Table of Contents


| Item | Title | Page |
|---|---|---|
| **Supplementary Methods** | | |
| | Hippocampal volumetry and definition of hippocampal atrophy | 3 |
| | Sensitivity analyses | |
| | PET normalisation sensitivity analysis | 4 |
| | Electroclinical measures operationalisation sensitivity analysis | 4 |
| | Partial-volume correction | 4 |
| **Supplementary Tables** | | |
| **Table 1** | Sensitivity analyses of the association between hippocampal $^{18}$F-MK6240 SUVR and memory performance | 5 |
| **Table 2** | Associations between ASM burden, hippocampal $^{18}$F-MK6240 SUVR, memory performance, and sleep microstructure | 5 |
| **Table 3** | Sensitivity analyses of hippocampal $^{18}$F-MK6240 SUVR in relation to electroclinical and radiological measures | 6 |
| **Table 4** | Associations of regional $^{18}$F-MK6240 uptake (Z-scores) in the temporal epileptogenic zone with seizure burden, NREM sleep microstructure, and memory performance | 7 |
| **Table 5** | Influence diagnostics and sensitivity analysis for the association between hippocampal $^{18}$F-M K6240uptake and N3 slow-wave rate | 8 |
| **Table 6** | Sensitivity analysis of hippocampal $^{18}$F-MK6240 SUVR after iterative Yang partial-volume correction | 8 |
| **Supplementary Figures** | | |
| **Figure 1** | Study flowchart | 9 |
| **Figure 2** | Peri-hippocampal and lateral temporal regional $^{18}$F-MK6240 SUVR and seizure burden measures | 10 |
| **Figure 3** | Peri-hippocampal and lateral temporal regional $^{18}$F-MK6240 SUVR, NREM sleep microstructure, and memory performance | 11 |
| **Figure 4** | Normal Q–Q plot of residuals | 12 |
| **Figure 5** | Residuals versus fitted values plot | 13 |
| **References** | | 14 |

### Supplementary Methods

##### Hippocampal volumetry and definition of hippocampal atrophy

Hippocampal volumes were derived from 3T T1-weighted MRI using the validated HippUnfold segmentation pipeline (1, 2), consistent with established approaches in temporal lobe epilepsy (TLE) imaging. Volumes were extracted for both hippocampi, adjusted for intracranial volume, and standardised relative to an age-matched healthy control cohort acquired on the same scanner using an identical imaging protocol, in line with standard normative transformation procedures commonly applied in temporal lobe epilepsy volumetry studies (3). Z-scores were defined as: (patient hippocampal volume − control mean hippocampal volume) / control standard deviation. Hippocampal asymmetry was assessed using hippocampal surface maps of cortical thickness derived from the same pipeline. For each participant, left- and right-hemisphere mean values were extracted across the hippocampal surface, and an asymmetry index was calculated as (LH − RH)/ [(LH + RH)/2]. Hippocampal atrophy was defined a priori as MRI-derived hippocampal volume or hippocampal asymmetry exceeding 1.5 standard deviations from the corresponding distribution of age-matched healthy controls (4). Quantitative classifications were cross-checked against the corresponding clinical neuroradiology reports.

#### Sensitivity analyses

##### PET normalisation sensitivity analysis

To assess whether findings were robust to the method of PET signal normalisation, all analyses were repeated across the three predefined regions of interest (hippocampus, peri-hippocampal regions, and lateral temporal regions) within the epileptogenic hemisphere using z-score–standardized PET measures.

Regional SUVR values were standardized relative to a cohort of 25 age- and sex-matched healthy controls who underwent comparable 3T MRI and $^{18}$F-MK6240 PET protocols (8 females; mean [SD] age, 34 [10·3] years; range, 24–60 years). Z-scores represented the deviation of individual regional uptake values from the corresponding normative distribution and were used to evaluate the consistency of findings across PET normalisation approaches, consistent with prior neuroimaging approaches (5).

##### Sensitivity analyses of electroclinical measures and hippocampal $^{18}$F-MK6240 SUVR

In addition to the prespecified categorical seizure burden classification, complementary sensitivity analyses were performed to assess the robustness of associations with hippocampal $^{18}$F-MK6240 uptake.

First, focal seizure frequency (events per month) was analysed as a continuous variable in relation to hippocampal $^{18}$F-MK6240 uptake, N2 spindle rate, N3 slow-wave rate, and memory performance using Pearson correlation analyses. Second, memory performance was analysed as a continuous variable in relation to hippocampal $^{18}$F-MK6240 uptake, N2 spindle rate, and N3 slow-wave rate using Pearson correlation analyses. Third, associations between continuous ipsilateral hippocampal volume, $^{18}$F-MK6240 uptake, and memory performance were examined. Finally, focal to bilateral tonic–clonic seizures (FBTCs) were analysed using an alternative binary classification (history present versus absent), irrespective of annual seizure frequency. Hippocampal $^{18}$F-MK6240 uptake was compared between groups using the Mann–Whitney U test because of non-normal data distribution.

Where regression models were performed, linearity, approximate normality of residuals, and homoscedasticity were assessed by inspection of scatterplots, residual Q–Q plots, and residual-versus-fitted plots, respectively. Potential influential observations were assessed using leverage and Cook's distance. Two observations with relatively high leverage were identified for the association between hippocampal $^{18}$F-MK6240 uptake and N3 slow-wave rate. Both observations were retained in the primary analysis because there was no prespecified clinical or methodological criterion for exclusion. A sensitivity analysis excluding these two observations was performed. The primary and sensitivity analyses were compared using effect estimates, 95% confidence intervals, and two-sided P values. Given the exploratory nature of these sensitivity analyses and the limited sample size, findings were interpreted primarily on the basis of effect estimates and 95% confidence intervals rather than P values alone.

##### Partial-volume correction

As a sensitivity analysis, PET images were additionally processed using iterative Yang partial-volume correction implemented in PETPVC (6, 7). The iterative Yang method uses anatomically defined regions to estimate regional activity and correct for spill-in and spill-out between regions. The method assumes approximate uniformity of tracer activity within the regions used to derive the correction factors and depends on accurate anatomical segmentation and PET–MRI registration, as well as appropriate characterisation of the scanner point-spread function.

Partial-volume correction was performed using 10 iterations, a 2.4 mm FWHM point-spread function, and FreeSurfer v6·0 anatomical segmentation. Partial-volume-corrected hippocampal $^{18}$F-MK6240 SUVR values were derived using the same cerebellar grey-matter reference region as in the primary analyses. The principal hippocampal analyses were repeated using these values.

#### Supplementary Tables

**Supplementary Table 1. Sensitivity analyses of associations between hippocampal $^{18}$F-MK6240 SUVR and memory performance**

**Part A: Binary memory impairment**

| Analysis | Effect estimate | 95% CI | P value |
|---|---|---|---|
| **Unadjusted group comparison** | 1·15 | 0·07 to 2·21 | 0·04 |
| **Hippocampal volume included** | $8·74 \times 10^{-5}$ | 0·00 to 3·13 | 0·07 |
| **Seizure-focus laterality included** | $1·86 \times 10^{-6}$ | 0·00 to 1·44 | 0·06 |

**Part B: Continuous memory performance**

| Analysis | Effect estimate | 95% CI | P value |
|---|---|---|---|
| **Unadjusted model** | −0·21 | −0·59 to 0·24 | 0·35 |
| **Hippocampal volume included** | −0·03 | −0·09 to 0·04 | 0·39 |
| **Seizure-focus laterality included** | −0·03 | −0·10 to 0·04 | 0·35 |

**Note: Part A** presents sensitivity analyses of the association between hippocampal MK-6240 SUVR and binary memory impairment, defined as a composite memory z score less than −1·3. The unadjusted analysis used an independent-samples t test, with Cohen's d reported as the effect estimate. Exploratory sensitivity analyses used binary logistic regression, with odds ratios (ORs) representing the change in odds associated with a 1-unit increase in the continuous predictor as entered into the model. Seizure-focus laterality was coded as right-sided versus left-sided focus. **Part B** presents analyses using continuous composite memory performance as the outcome. The unadjusted association was assessed using Pearson correlation, with the correlation coefficient (r) reported as the effect estimate. Sensitivity analyses adjusted for hippocampal volume or seizure-focus laterality used linear regression, with unstandardised regression coefficients (B) reported. Hippocampal $^{18}$F-MK6240 SUVR was the exposure of interest in all analyses. All P values are two-sided.

**Supplementary Table 2. Associations between ASM burden, hippocampal $^{18}$F-MK6240 SUVR, memory performance, and sleep microstructure**

| Measure | N | Effect estimate | 95% CI | P value |
|---|---|---|---|---|
| **ASM burden** | | | | |
| **$^{18}$F-MK6240 SUVR** | 25 | −0·22 | −0·57 to 0·19 | 0·29 |
| **Continuous memory performance** | 21 | 0·26 | −0·20 to 0·62 | 0·26 |
| **Memory impairment (impaired vs intact)** | 21 | −0·33 | −1·33 to 0·69 | 0·53 |
| **N3 slow-wave rate** | 25 | −0·07 | −0·45 to 0·34 | 0·74 |
| **N2 spindle rate** | 25 | 0·65 | 0·34 to 0·83 | <0·001 |
| **N2 spindle rate (adjusted for ASM burden) → $^{18}$F-MK6240 SUVR** | 25 | −0·05 | −0·11 to 0·01 | 0·08 |

**Note:** Pearson correlation coefficients were used for associations between continuous variables. Cohen’s d represents the standardized mean difference in ASM burden between participants with impaired and intact memory performance. The adjusted analysis used linear regression with hippocampal $^{18}$F-MK6240 SUVR as the outcome and N2 spindle rate as the predictor, adjusting for ASM burden. All P values are two-sided and unadjusted.

**Supplementary Table 3. Sensitivity analyses of hippocampal $^{18}$F-MK6240 SUVR in relation to electroclinical and radiological measures**

| Electroclinical measures | Outcome | N | Summary measure | Effect estimate | 95% CI | p value |
|---|---|---|---|---|---|---|
| **Seizure burden** | | | | | | |
| **Focal seizures (per month)** | $^{18}$F-MK6240 SUVR | 25 | — | 0·22 | −0·19 to 0·57 | 0·28 |
| | N2 spindle rate | 25 | — | 0·28 | −0·13 to 0·61 | 0·17 |
| | N3 slow-wave rate | 25 | — | 0·05 | −0·36 to 0·43 | 0·83 |
| | Memory performance | 21 | — | 0·13 | −0·32 to 0·53 | 0·58 |
| **FTBCs (present; n=19 vs absent; n=6)** | $^{18}$F-MK6240 SUVR | 25 | Median [IQR]: 0·67 [0·62–0·75] vs 0·69 [0·66–0·76] | −0·09 | −0·56 to 0·42 | 0·78 |
| **Memory performance** | | | | | | |
| | $^{18}$F-MK6240 SUVR | 21 | — | −0·21 | −0·59 to 0·24 | 0·35 |
| | N2 spindle rate | 21 | — | 0·29 | −0·16 to 0·64 | 0·20 |
| | N3 slow-wave rate | 21 | — | 0·01 | −0·42 to 0·44 | 0·97 |
| **Hippocampal volume** | | | | | | |
| | $^{18}$F-MK6240 SUVR | 25 | — | −0·20 | −0·55 to 0·21 | 0·34 |
| | Memory performance | 21 | — | −0·06 | −0·38 to 0·48 | 0·79 |
| **Radiological MR findings (MTS; n =5 vs others; n = 20)** | $^{18}$F-MK6240 SUVR | 25 | Mean [SD]: 0·75 [0·12] vs 0·69 [0·11] | 0·53 | −0·47 to 1·51 | 0·30 |

**Abbreviations:** 95% CI, confidence interval; FBTCs, focal to bilateral tonic–clonic seizures; $^{18}$F-MK6240, $^{18}$F-MK6240 tau PET tracer; MTS, mesial temporal sclerosis; NREM, non-rapid eye movement; N2/N3, NREM sleep stages N2 and N3. **Note:** Effect estimates are presented as Pearson correlation coefficients, rank-biserial correlation, or Cohen's d, as appropriate, with corresponding 95% confidence intervals. All P values are two-sided and unadjusted.

**Supplementary Table 4. Associations of Regional [18]F-MK6240 Uptake (Z-scores) in the Temporal Epileptogenic Zone with Seizure Burden, NREM Sleep Microstructure, and Memory Performance**

| ROI | Domain | Electroclinical measures | Summary measure | Effect estimate | 95% CI | P value |
|---|---|---|---|---|---|---|
| **Hippocampus** | | | | | | |
| | **Seizure burden** | Focal seizure burden (≥1/week; high; n=16 vs low; n=9) | Median [IQR]: 0·67 [0·64–0·74] vs 0·64 [0·58–0·65] | −0·58 | −0·82 to −0·19 | 0·02 |
| | | FBTCs burden (≥3/year; high; n=8 vs low; n=17) | Mean [SD]: 0·13 [1·22] vs −0·24 [0·79] | 0·38 | −0·47 to 1·23 | 0·38 |
| | | IED frequency (≥1 event/min; frequent; n=15 vs non-frequent; n=10) | Mean [SD]: 0·09 [0·91] vs −0·44 [0·94] | 0·58 | −0·25 to 1·39 | 0·17 |
| | **NREM sleep microstructure** | N2 spindle rate (events/min; n=25) | — | −0·42 | −0·70 to −0·03 | 0·04 |
| | | N3 slow-wave rate (events/min; n=25) | — | 0·47 | 0·09 to 0·73 | 0·02 |
| | **Memory performance** | Impaired (n=5) vs intact (n=16) | Mean [SD]: 0·74 [1·05] vs −0·30 [0·85] | 1·15 | 0·07 to 2·21 | 0·04 |
| **Peri-hippocampal temporal region** | | | | | | |
| | **Seizure burden** | High focal seizure frequency (≥1/week; high; n=16 vs low; n=9) | Mean [SD]: −0·65 [0·82] vs 0·10 [1·01] | 0·79 | -0·07 to 1·63 | 0·07 |
| | | FBTCs burden (≥3/year; high; n=8 vs low; n=17) | Mean [SD]: −0·15 [0·91] vs −0·18 [1·06] | 0·03 | −0·81 to 0·87 | 0·95 |
| | | IED frequency (≥1 event/min; frequent; n=15 vs non-frequent; n=10) | Mean [SD]: −0·09 [1·02] vs −0·28 [1·01] | 0·18 | −0·62 to 0·98 | 0·66 |
| | **NREM sleep microstructure** | N2 spindle rate (events/min; n=25) | — | −0·24 | −0·58 to 0·18 | 0·26 |
| | | N3 slow-wave rate (events/min; n=25) | — | 0·23 | −0·18 to 0·57 | 0·27 |
| | **Memory performance** | Impaired (n=5) vs intact (n=16) | Mean [SD]: 0·39 [1·00] vs −0·31 [0·98] | 0·71 | −0·33 to 1·73 | 0·18 |
| **Lateral temporal region** | | | | | | |
| | **Seizure burden** | High focal seizure frequency (≥1/week; high; n=16 vs low; n=9) | Mean [SD]: −0·25 [1·45] vs 0·48 [1·00] | 0·54 | -0·30 to 1·37 | 0·21 |
| | | FBTCs burden (≥3/year; high; n=8 vs low; n=17) | Mean [SD]: 0·16 [0·65] vs 0·28 [1·69] | −0·08 | −0·92 to 0·76 | 0·85 |
| | | IED frequency (≥1 event/min; frequent; n=15 vs non-frequent; n=10) | Mean [SD]: 0·40 [1·52] vs 0·00 [1·33] | 0·28 | −0·53 to 0·98 | 0·51 |
| | **NREM sleep microstructure** | N2 spindle rate (events/min; n=25) | — | −0·03 | −0·42to 0·37 | 0·87 |
| | | N3 slow-wave rate (events/min; n=25) | — | 0·08 | −0·33 to 0·46 | 0·72 |
| | **Memory performance** | Impaired (n=5) vs intact (n=16) | Mean [SD]: 0·12 [1·76] vs 0·07 [1·41] | 0·03 | −0·98 to 1·03 | 0·96 |

**Abbreviations:** 95% CI, confidence interval; FBTCs, focal to bilateral tonic–clonic seizures; [18]F-MK6240, tau PET tracer; IED, interictal epileptiform discharges; NREM, non–rapid eye movement; N2/N3, NREM sleep stages N2 and N3; ROI, region of interest; SD, standard deviation. **Note:** Between-group comparisons are expressed as mean (SD) or median (IQR). Effect sizes are reported as Cohen d for parametric comparisons, rank-biserial correlation for nonparametric comparisons, and Pearson correlation coefficient for associations. All confidence intervals are 95% and are reported as lower and upper bounds. All P values are two-sided and unadjusted.

**Supplementary Table 5. Influence diagnostics and sensitivity analysis for the association between hippocampal $^{18}$F-MK6240 uptake and N3 slow-wave rate**

- **Part A: Influence diagnostics**

| Observation | Leverage | Cook's distance |
|---|---|---|
| **Case 7** | 0·251 | 0·082 |
| **Case 20** | 0·271 | 0·119 |

**Note:** These were the two observations identified as having relatively high leverage. Their Cook's distances were low, indicating limited individual influence on the fitted association. Both observations were retained in the primary analysis.

- **Part B: Sensitivity analysis**

| Analysis | n | Pearson r | 95% CI | P value |
|---|---|---|---|---|
| **Primary analysis** | 25 | 0·47 | 0·09 to 0·73 | 0·02 |
| **Excluding high-leverage observations** | 23 | 0·38 | −0·04 to 0·68 | 0·08 |

**Note:** Exclusion of the two high-leverage observations did not materially alter the direction or magnitude of the estimated association, although the confidence interval became wider because of the reduced sample size.

- **Supplementary Table 6: Sensitivity analysis of hippocampal $^{18}$F-MK6240 SUVR after iterative Yang partial-volume correction**

| Electroclinical Measure | Outcome | N | Effect estimate | 95% CI | p value |
|---|---|---|---|---|---|
| **Seizure burden** | | | | | |
| Focal seizures (≥1/week vs <1/week) | $^{18}$F-MK6240 | 25 | −0·49 | −0·77 to −0·05 | 0·049 |
| FBTCs (≥3/year vs <3/year) | $^{18}$F-MK6240 | 25 | 0·00 | −0·46 to 0·46 | 0·895 |
| **NREM sleep microstructure** | | | | | |
| N2 spindle rate | $^{18}$F-MK6240 | 25 | −0·38 | −0·67 to 0·02 | 0·06 |
| N3 slow-wave rate | $^{18}$F-MK6240 | 25 | 0·50 | 0·14 to 0·75 | 0·01 |
| **Memory performance** | | | | | |
| Memory impairment (impaired vs intact) | $^{18}$F-MK6240 | 21 | 1·09 | 0·02 to 2·14 | 0·046 |

**Abbreviations:** CI, confidence interval; FBTCs, focal to bilateral tonic–clonic seizures; $^{18}$F-MK6240, tau PET tracer; NREM, non-rapid eye movement; N2/N3, NREM sleep stages N2 and N3; SUVR, standardised uptake value ratio. **Note:** Analyses were repeated using partial-volume-corrected ipsilateral hippocampal $^{18}$F-MK6240 SUVR values derived using the iterative Yang method. Effect estimates and corresponding 95% CIs are reported according to the statistical tests used in the primary analyses. All p values are two-sided and unadjusted.

#### Supplementary Figures

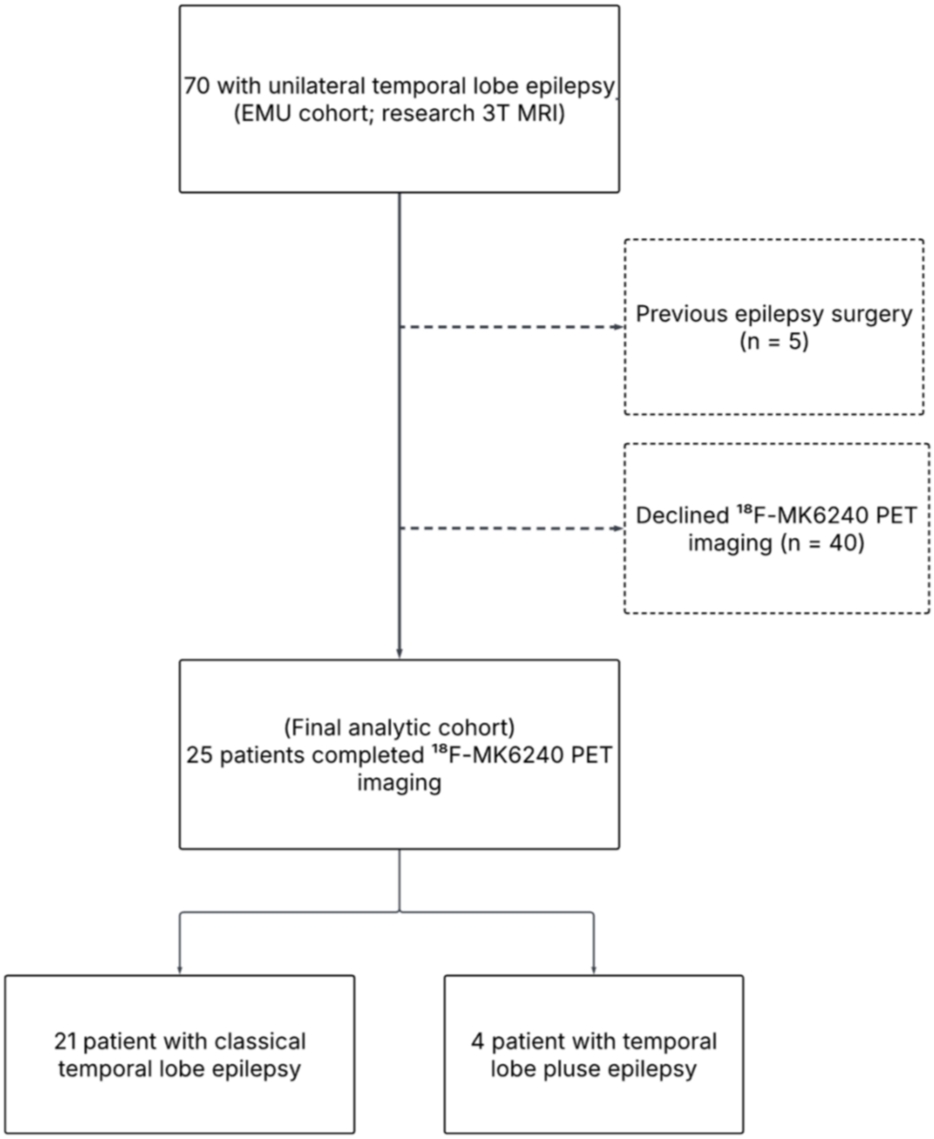


**Supplementary Figure 1. Study flowchart.**

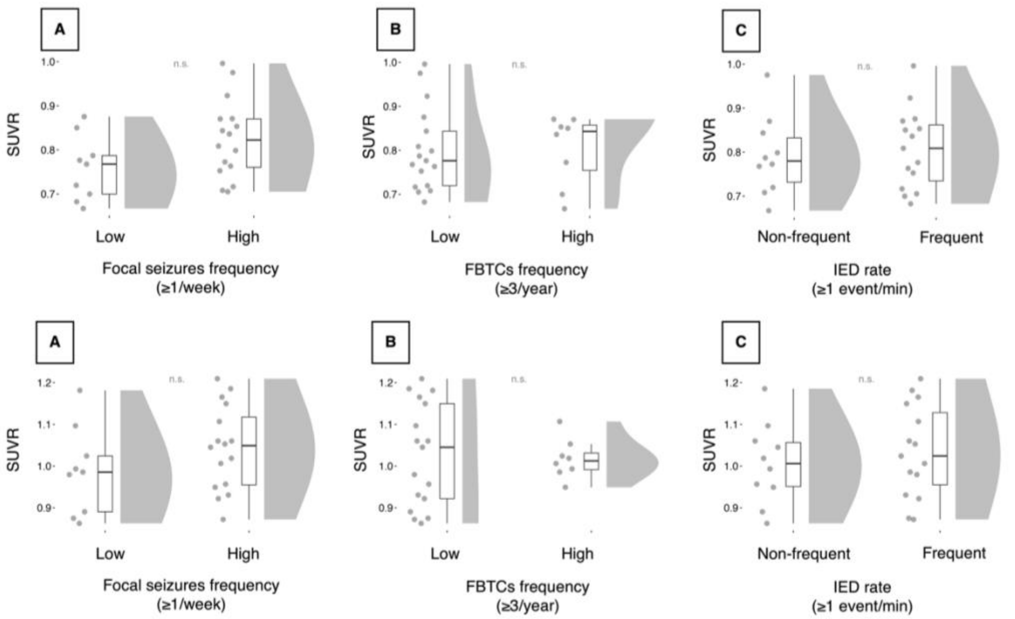


**Supplementary Figure 2. Peri-hippocampal and lateral temporal regional $^{18}$F-MK6240 SUVR and seizure burden measures.**
**Top row – Peri-hippocampal temporal regions.**
(A) Peri-hippocampal $^{18}$F-MK6240 SUVR and focal seizure burden (high vs low burden; ≥1 seizure/week; n=16 vs n=9) (mean [SD] 0·83 [0·09] vs 0·76 [0·07], Cohen's d=0·79, 95% CI −0·07 to 1·63; p=0·07).
(B) Peri-hippocampal $^{18}$F-MK6240 SUVR and focal to bilateral tonic–clonic seizure burden (FBTCs ≥3/year; high vs low burden; n=8 vs n=17) (mean [SD] 0·80 [0·08] vs 0·80 [0·09], Cohen's d=0·03, 95% CI −0·81 to 0·87; p=0·95).
(C) Peri-hippocampal $^{18}$F-MK6240 SUVR and interictal epileptiform discharge burden (IED ≥1 event/min; frequent vs non-frequent; n=15 vs n=10) (mean [SD] 0·81 [0·09] vs 0·79 [0·09], Cohen's d=0·18, 95% CI −0·62 to 0·98; p=0·66).

**Bottom row – Lateral temporal regions.**
(A) Lateral temporal $^{18}$F-MK6240 SUVR and focal seizure burden (high vs low burden; ≥1 seizure/week; n=16 vs n=9) (mean [SD] 1·04 [0·10] vs 0·99 [0·11], Cohen's d=0·54, 95% CI −0·30 to 1·37; p=0·21).
(B) Lateral temporal $^{18}$F-MK6240 SUVR and focal to bilateral tonic–clonic seizure burden (FBTCs ≥3/year; high vs low burden; n=8 vs n=17) (mean [SD] 1·02 [0·05] vs 1·03 [0·12], Cohen's d=−0·08, 95% CI −0·92 to 0·76; p=0·85).
(C) Lateral temporal $^{18}$F-MK6240 SUVR and interictal epileptiform discharge burden (IED ≥1 event/min; frequent vs non-frequent; n=15 vs n=10) (mean [SD] 1·04 [0·11] vs 1·01 [0·10], Cohen's d=0·28, 95% CI −0·53 to 1·08; p=0·51).

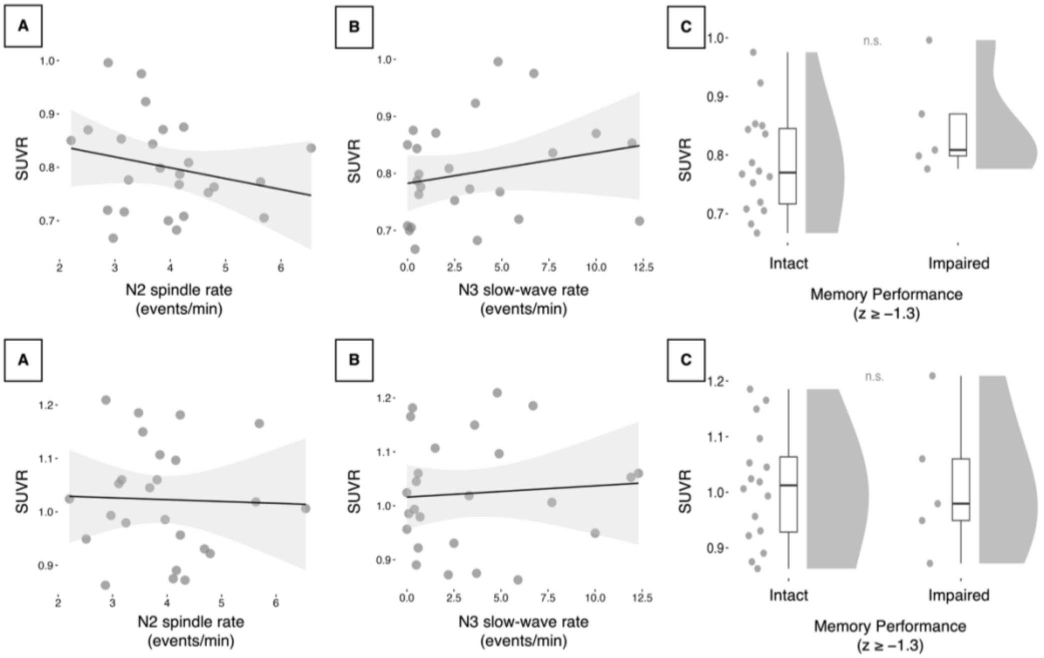


**Supplementary Figure 3. Peri-hippocampal and lateral temporal regional $^{18}$F-MK6240 SUVR, NREM sleep microstructure, and memory performance.**

**Top row – Peri-hippocampal temporal regions.**

(A) Peri-hippocampal $^{18}$F-MK6240 SUVR and N2 spindle rate (n=25; r=−0·24, 95% CI −0·58 to 0·18; p=0·26).

(B) Peri-hippocampal $^{18}$F-MK6240 SUVR and N3 slow-wave rate (n=25; r=0·23, 95% CI −0·18 to 0·57; p=0·27).

(C) Peri-hippocampal $^{18}$F-MK6240 SUVR in participants with impaired versus intact memory performance (n=5 vs n=16; mean [SD] 0·85 [0·09] vs 0·79 [0·09], Cohen's d=0·71, 95% CI −0·33 to 1·73; p=0·18).

**Bottom row – Lateral temporal regions.**

**(A)** Lateral temporal $^{18}$F-MK6240 SUVR and N2 spindle rate (n=25; r=−0·03, 95% CI −0·42 to 0·37; p=0·87).

**(B)** Lateral temporal $^{18}$F-MK6240 SUVR and N3 slow-wave rate (n=25; r=0·08, 95% CI −0·33 to 0·46; p=0·72).

**(C)** Lateral temporal $^{18}$F-MK6240 SUVR in participants with impaired versus intact memory performance (n=5 vs n=16; mean [SD] 1·01 [0·13] vs 1·01 [0·10], Cohen's d=0·03, 95% CI −0·98 to 1·03; p=0·96).

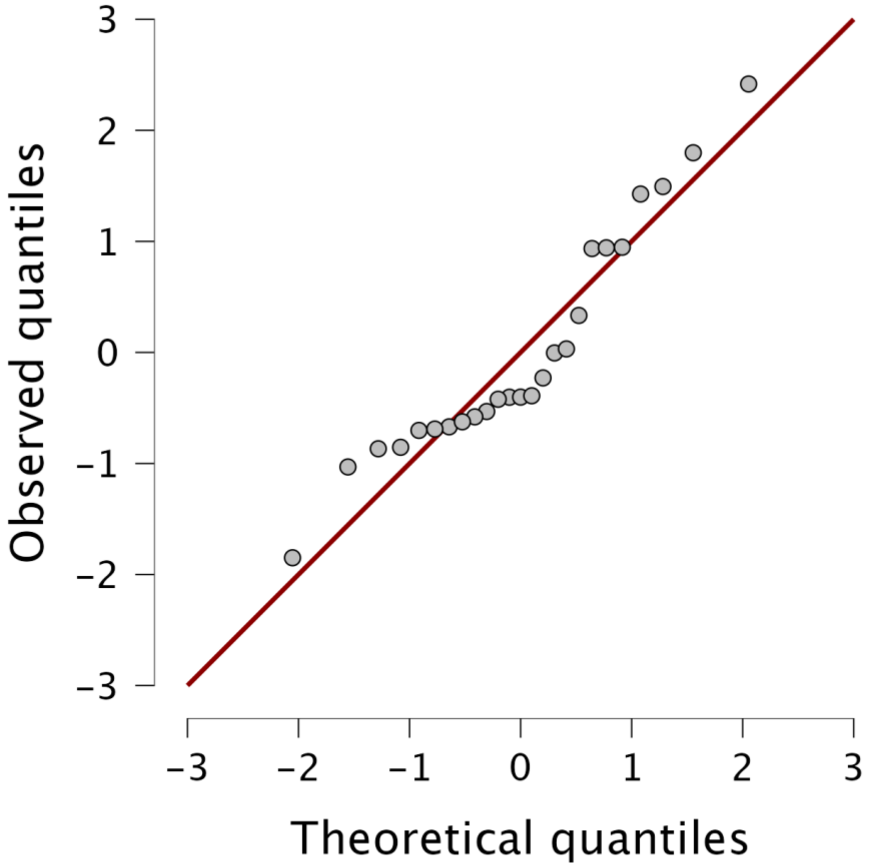


**Supplementary Figure 4: Normal Q–Q plot of residuals.** Residuals were approximately normally distributed, with only minor deviations from the reference line at the distribution tails.

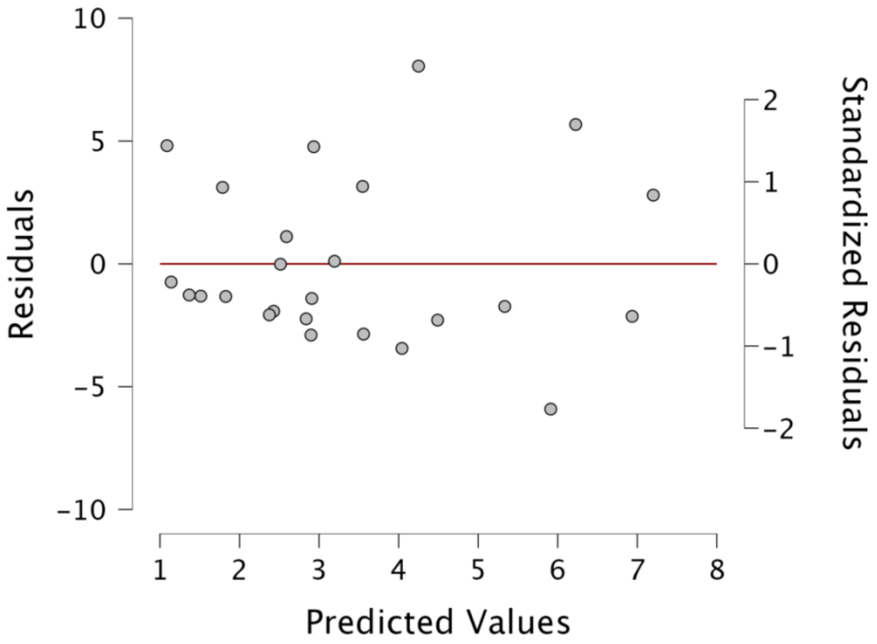


**Supplementary Figure 5: Residuals vs fitted values plot.** Visual inspection did not suggest substantial violations of the assumptions of linearity or homoscedasticity.
